# Evaluating sources of technical variability in the mechano-node-pore sensing pipeline and their effect on the reproducibility of single-cell mechanical phenotyping

**DOI:** 10.1101/2021.06.02.446242

**Authors:** Brian Li, Kristen L. Cotner, Nathaniel K. Liu, Stefan Hinz, Mark A. LaBarge, Lydia L. Sohn

**Author notes:** Corresponding author (LLS). These authors contributed equally to this work (Brian Li, Kristen L. Cotner).

## Abstract

Cellular mechanical properties can reveal physiologically relevant characteristics in many cell types, and several groups have developed microfluidics-based platforms to perform single-cell mechanical testing with high throughput. However, prior work has performed only limited characterization of these platforms’ technical variability and reproducibility. Here, we evaluate the repeatability performance of mechano-node-pore sensing, which is a single-cell mechanical phenotyping platform developed by our research group. We measured the degree to which device-to-device variability and semi-manual data processing affected this platform’s measurements of single-cell mechanical properties, and we demonstrated high repeatability across the entire technology pipeline even for novice users. We then compared results from identical mechano-node-pore sensing experiments performed by researchers in two different labs with different analytical instruments, demonstrating that the mechanical testing results from these two locations are in agreement. Our findings quantify the expectation of technical variability in mechano-node-pore sensing even in minimally experienced hands. Most importantly, we find that the repeatability performance we measured is fully sufficient for interpreting biologically relevant single-cell mechanical measurements with high confidence.

## Introduction

As cells frequently generate and experience a variety of forces in normal physiology, their mechanical properties are an important aspect of their function. Cell mechanical properties are implicated in many diseases including metastatic cancer and various laminopathies [1,2]. More recently, there has been increasing research on the use of “mechanical phenotyping” to identify and screen single cells for mechanical properties associated with malignancies [3–5]. In this work, we focus specifically on the use of mechano-node-pore sensing (mechano-NPS) for analyzing cells from breast epithelial tissue and from a drug-resistant leukemia cell line [5,6]. To extract mechanical information from cells, mechano-NPS uses a four-terminal measurement across a microchannel to track cell trajectories as they are deformed by a narrow constriction [7,8] (Fig 1A). Mechano-NPS, like other microfluidic methods for single-cell mechanical testing, has demonstrated enhanced deformability in cancer cells, reflecting invasive potential [1,3,5,9,10]. Uniquely, by measuring cell recovery from deformation, mechano-NPS has also been used to uncover age-dependent changes in viscoelastic properties and offers roughly 10-fold faster throughput for viscoelastic testing over older methods such as optical tweezers [5,11].

**Fig 1.**
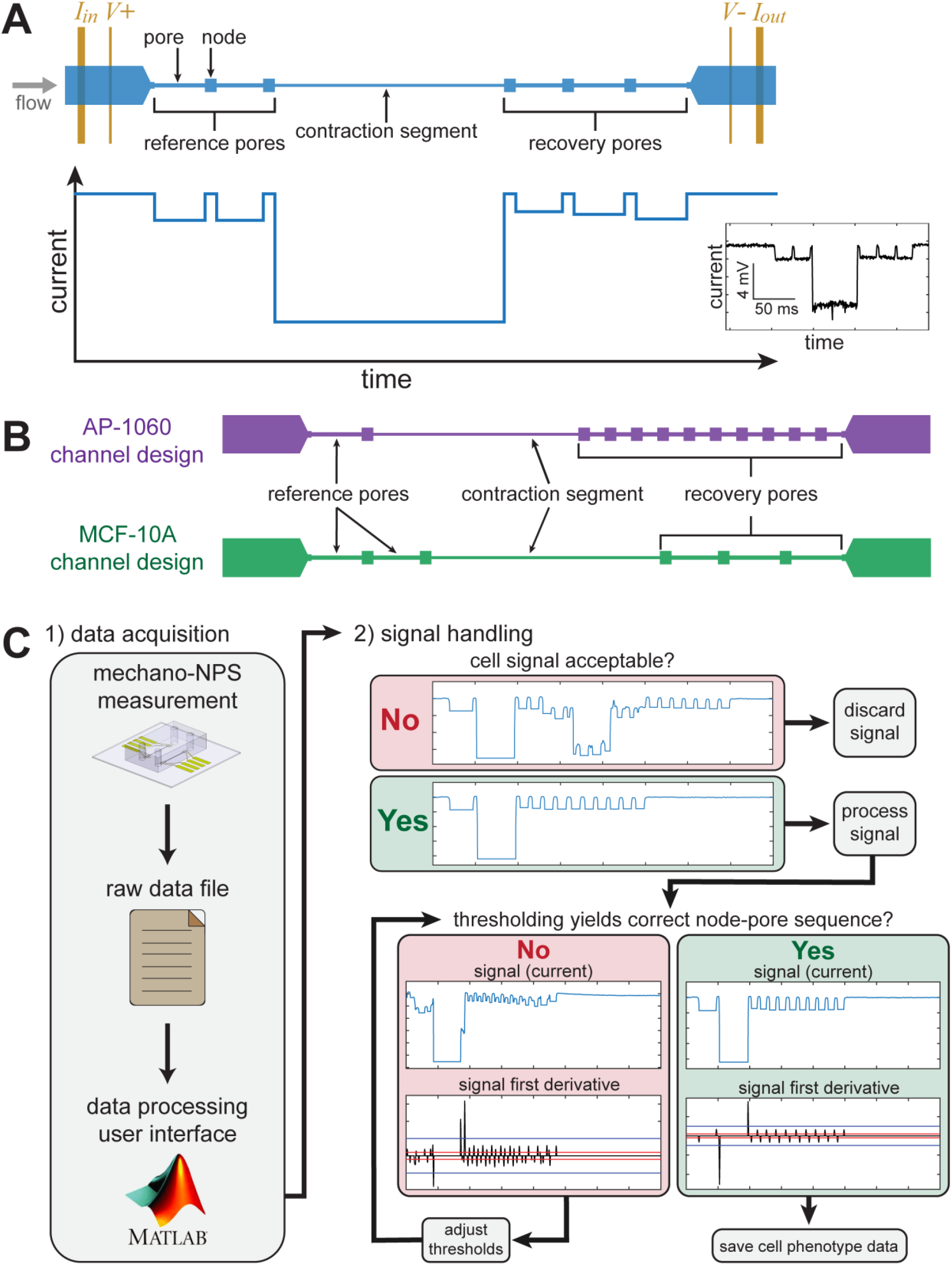
Overview of mechano-node-pore sensing (mechano-NPS) operating principles, device design, and data processing pipeline. (A) Top-down schematic view of a mechano-NPS device (top), with corresponding expected electric current pulse (bottom) caused by a single cell transiting the microfluidic channel. A potential is applied across the channel, causing a drop in measured current when a cell enters a narrow segment of the microchannel (pore). Inset: An actual current pulse caused by an MCF-10A cell traversing the channel. (B) Top-down schematic views of mechano-NPS channel designs used in this work for screening AP-1060 and MCF-10A cells. (C) Flow of user input to the data processing analysis software. The user must identify valid signals corresponding to cell measurements and exclude non-valid pulses. Subsequently, the user must set processing thresholds to the appropriate levels for each valid cell.

Previously, the technical variability of microfluidic techniques like mechano-NPS has only been addressed through simple validation and calibration. For mechano-NPS, Kim et al. made comparisons to published measurements of cortical tension and elastic modulus for several cell lines, showing that their measure of deformability followed published trends established with atomic force microscopy and micropipette aspiration [5]. Similarly, a comparison of the techniques employed by three different research groups demonstrated that these different technologies can, with limited agreement, sense similar trends in deformability [12]. To calibrate their technology known as hydrodynamic stretching, Gossett et al. carried out measurements on droplets of various viscosities, demonstrating the relationship between measured deformability and known values of viscosity [13]. Using suspended microchannel resonators, Kang et al. calibrated sensors to the mechanical properties of polystyrene beads and hydrogel spheres of varying elastic modulus [14]. For real-time deformability cytometry, theoretical modeling of soft matter deformation was additionally tested against agar and polyacrylamide beads of varying stiffnesses to validate observed changes in apparent cell deformability [15,16]. Notably, only two previous works (Gossett et al. and Kim et al.) performed any reliability testing of their respective platforms, with both works assessing only a handful of sources of variability [5,13]. Thus, it is clear that the literature is lacking in reproducibility analyses for microfluidic platforms that measure single-cell mechanical properties. Such analysis provides a critical performance benchmark for other researchers who may seek to adopt these technologies.

Observing this need for reproducibility analyses in our field, we set out to examine the reproducibility of the mechano-NPS system and its measurements of single-cell mechanical phenotypes. First, to characterize device-to-device variability, we quantified the differences in mechano-NPS results when a single biological sample is tested across multiple replicate devices with relatively low sample sizes. Next, we examined the intra- and inter-user reliability of the semi-manual mechano-NPS data processing pipeline by analyzing results obtained by several researchers using our MATLAB command-line interface (CLI) [6] to process identical mechano-NPS raw data sets. Last, to assess the reproducibility of the mechano-NPS technology platform, we evaluated the similarity in results from two identical experiments conducted on two different sets of mechano-NPS hardware by different researchers in different locations.

Overall, we show that average measures of single-cell mechanical parameters using mechano-NPS are highly repeatable. Using our new CLI, both experienced and novice users were able to rapidly process hundreds of cell events and produce highly consistent results. We show that the current mechano-NPS pipeline is capable of high degrees of consistency, high throughput (allowing for large sample sizes), and rapid analysis of large data sets. This work provides an important insight into the performance and reproducibility of mechano-NPS, demonstrating its maturity as a technology and potential for further adoption by other research groups.

## Results and discussion

### Mechano-NPS principles, device design, and testing procedures

In mechano-NPS, as cells flow through a microfluidic channel, the channel’s electrical resistance is measured using a four-terminal measurement (Fig 1A). The channel is segmented into wider “nodes” and narrower “pores” that generate resistive subpulses which, together, comprise a unique resistive pulse when a cell transits the channel (Fig 1A). The cell first passes through reference pore(s) in order to measure its original size and velocity, then passes through a narrow contraction segment designed to target a specific strain (percent-change in cell diameter along the axis of deformation) and measure its resistance to deformation. After passing through the contraction segment, the cell transits several recovery pores in which its recovery from the deformation applied in the contraction segment is measured as it returns to its original size.

The geometry of each mechano-NPS device is optimized to measure a particular cell size. To assess the reproducibility of previous studies using mechano-NPS, we emulated devices designed by Kim et al. and Li et al. to measure MCF-10A and AP-1060 cells, respectively [5,6] (Fig 1B). Both devices assess cell elastic deformability according to a cell’s transit time through the contraction segment using a unitless number called the whole-cell deformability index (*wCDI*), which is inversely related to a cell’s Young’s modulus [5]. Cell viscoelastic behavior is assessed by measuring its recovery from deformation. For MCF-10A cells measured with the device design described by Kim et al., cells are classified into distinct recovery categories defined by how long the cell took to return to its original shape after deformation. For AP-1060 cells measured with the device design described by Li et al., a quantitative recovery time constant (*τ*) is calculated from the rate at which the cell relaxes from a deformed ellipsoid to a sphere. We first evaluated how device-to-device variability and our data processing pipeline (Fig 1C) affect mechano-NPS reproducibility, using AP-1060 cells as an example. We then assessed the overall reproducibility of mechano-NPS measurements in two different laboratories using identical devices and identical MCF-10A cells.

### Device-to-device variability and its effects on mechanical phenotyping

We first sought to characterize how device-to-device variability could affect mechano-NPS reproducibility, which can be masked when pooling data from several replicate devices and samples. As such, we used mechano-NPS to mechanically phenotype relatively small samples (46 ≤ *n* ≤ 79) of AP-1060 cells, using three different devices. Using the mechano-NPS device design described by Li et al. [6] (Fig 1B), we measured the *wCDI* and recovery time constant *τ* of all AP-1060 samples.

We tested each of the *wCDI* distributions (Fig 2A) collected from Devices 1, 2, and 3 for normality and found only Device 3 to be non-normal (*p =* 0.016) (S1 Table). We evaluated whether the data from these three groups were sampled from the same distribution with a Kruskal-Wallis test and found that these data reject the null hypothesis of an equal originating distribution (*p =* 0.014). Pairwise comparisons for all three devices showed that Device 1 and Device 3 were not significantly different (*p =* 0.72), while Device 2 was statistically significantly different from Device 1 (*p =* 0.005) and Device 3 (*p =* 0.009) (Fig 2A). These results, along with analysis of the empirical distribution functions (CDFs) (Fig 2A), clearly show that Devices 1 and 3 produced highly similar *wCDI* results, whereas *wCDI* data from Device 2 were significantly lower. Pairwise tests of *wCDI* CDFs found that while Device 1 and 2 *wCDI* distributions were not significantly different (*p*_1,2_ = 0.024, *α* = 0.0167), Device 2 and Device 3 data were not sampled from the same distribution, with Device 2 data again being significantly lower (*p*_2,3_ = 0.012, *α* = 0.0167).

**Fig 2.**
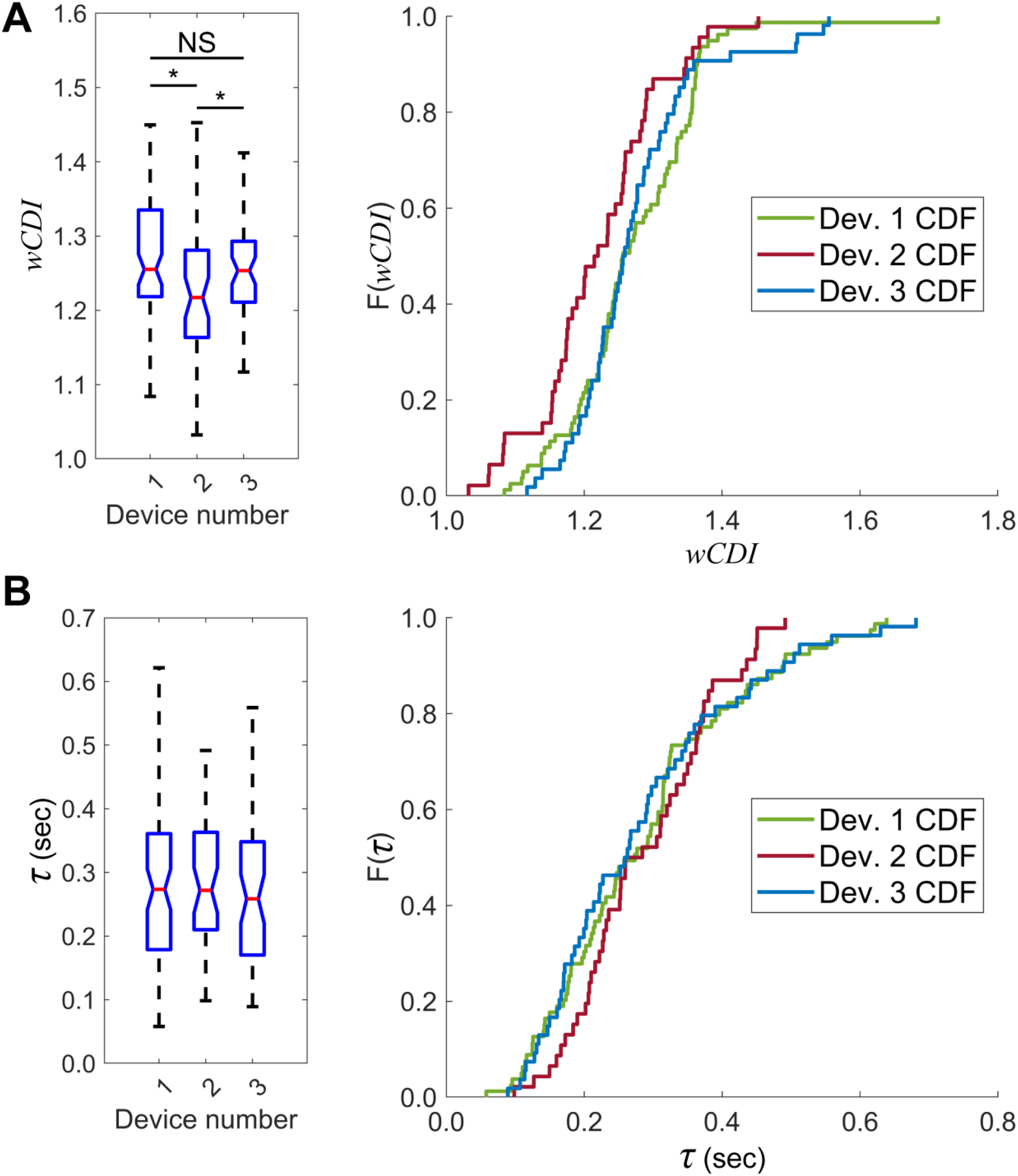
Between-device variability of mechanical phenotyping parameters measured by mechano-NPS. (A) Left: Box plots of whole-cell deformability index (*wCDI*) for a single sample of AP-1060 cells analyzed using three separate devices. Notches represent 95% confidence intervals for the true sample median for each distribution. * indicates *p <* 0.0167; NS indicates no statistical significance. Statistical significance between devices was determined using pairwise Wilcoxon rank-sum tests with a Bonferroni correction. (*p*_1,2_ = 0.005, *p*_1,3_ = 0.72, *p*_2,3_ = 0.009; *n*_1_ = 79, *n*_2_ = 46, *n*_3_ = 53; tested against a Bonferroni-corrected significance criterion of 0.05: *α =* 0.0167). Right: Empirical cumulative distribution functions (CDF) for *wCDI* data. Pairwise two-sample Kolmogorov-Smirnov tests determined that only Device 2 and Device 3 data were sampled from different distributions (*p*1,2 = 0.024, *p*1,3 = 0.55, *p*2,3 = 0.012; *n*1 = 79, *n*2 = 46, *n*3 = 53; tested against a Bonferroni-corrected significance criterion of 0.05: *α =* 0.0167). (B) Box plots of recovery time constant (*τ*) for the AP-1060 cells analyzed in (A) on each of three devices. Notches represent 95% confidence intervals for the true sample median for each distribution. A Kruskal-Wallis test failed to reject the null hypothesis that the recovery time constants measured by each device came from the same distribution. *n*_1_ = 79, *n*_2_ = 46, *n*_3_ = 53. Right: Empirical CDFs for recovery time constant data.

We performed a similar analysis on the distributions of recovery time constant (Fig 2B), first identifying that the distribution from Device 1 was non-normal (*p =* 0.009). Analyzing the recovery data obtained from these three devices, we found that the recovery time constant data for all three groups were sampled from the same distribution (*p =* 0.52), precluding the need for further pairwise comparisons. This analysis suggests that compared to *wCDI*, the measurement of continuous recovery time constant is less sensitive to device-to-device variability and is thus more robust against variability in manufacturing.

For mechano-NPS, the data from replicate devices are pooled before comparisons are made between experimental conditions [5,6]. The discrepancy among *wCDI* data from the different devices highlights the importance of this data pooling, which reduces the influence of device-to-device variability. Even though similar variation was not observed in recovery time constant measurements, data pooling would serve the same purpose. In comparing the difference in median *wCDI* among the three devices, there was only a 3.25% difference in median between Device 2 and Device 1 or 3. While biological variability may also influence this result, each of the three samples of AP-1060 cells were biological replicates and were handled identically prior to mechano-NPS measurements (see Methods). While this served to minimize differences between replicates, biological variability is impossible to completely eliminate. As such, the measured 3.25% difference in median *wCDI* may be an overestimate of the actual device-to-device variability in mechano-NPS. Overall, this experiment quantifies the degree to which device-to-device variability in mechano-NPS can affect measurements of *wCDI* and recovery time constant.

### Intra- and inter-user reliability of results obtained using mechano-NPS data processing pipeline

As the processing of mechano-NPS data is another critical aspect for producing consistent results, we quantified the variability introduced through the semi-manual nature of our custom mechano-NPS data processing software [6] (Fig 1C). We recruited five subjects who are microfluidics researchers with varying degrees of familiarity with mechano-NPS and surveyed them about their familiarity with the mechano-NPS data processing software. Subjects 1–2 identified as experienced users of the software, while Subjects 3–5 identified as novice users of the software. We then provided all subjects with a copy of the mechano-NPS data processing software and a 30-minute training video on how to use it. All subjects were tasked with performing data processing on five blinded mechano-NPS raw data sets using the custom software. These data sets comprised five measurements (A–E) of AP-1060 cells (Fig 3). Data sets A–C were obtained from measuring untreated cells, while data sets D–E were obtained from measuring cells treated with the microfilament disrupting agent LatrunculinA (LatA). Additionally, each subject performed analysis on each data set three different times in order to test intra-user reliability. Overall, we examined both intra- and inter-user reliability using the results from experienced and novice users of the software. This human subjects work was conducted under an IRB-approved protocol (UC Berkeley Committee for Protection of Human Subjects, Protocol ID: 2020-11-13822), and all subjects provided written informed consent prior to taking part in the study.

**Fig 3.**
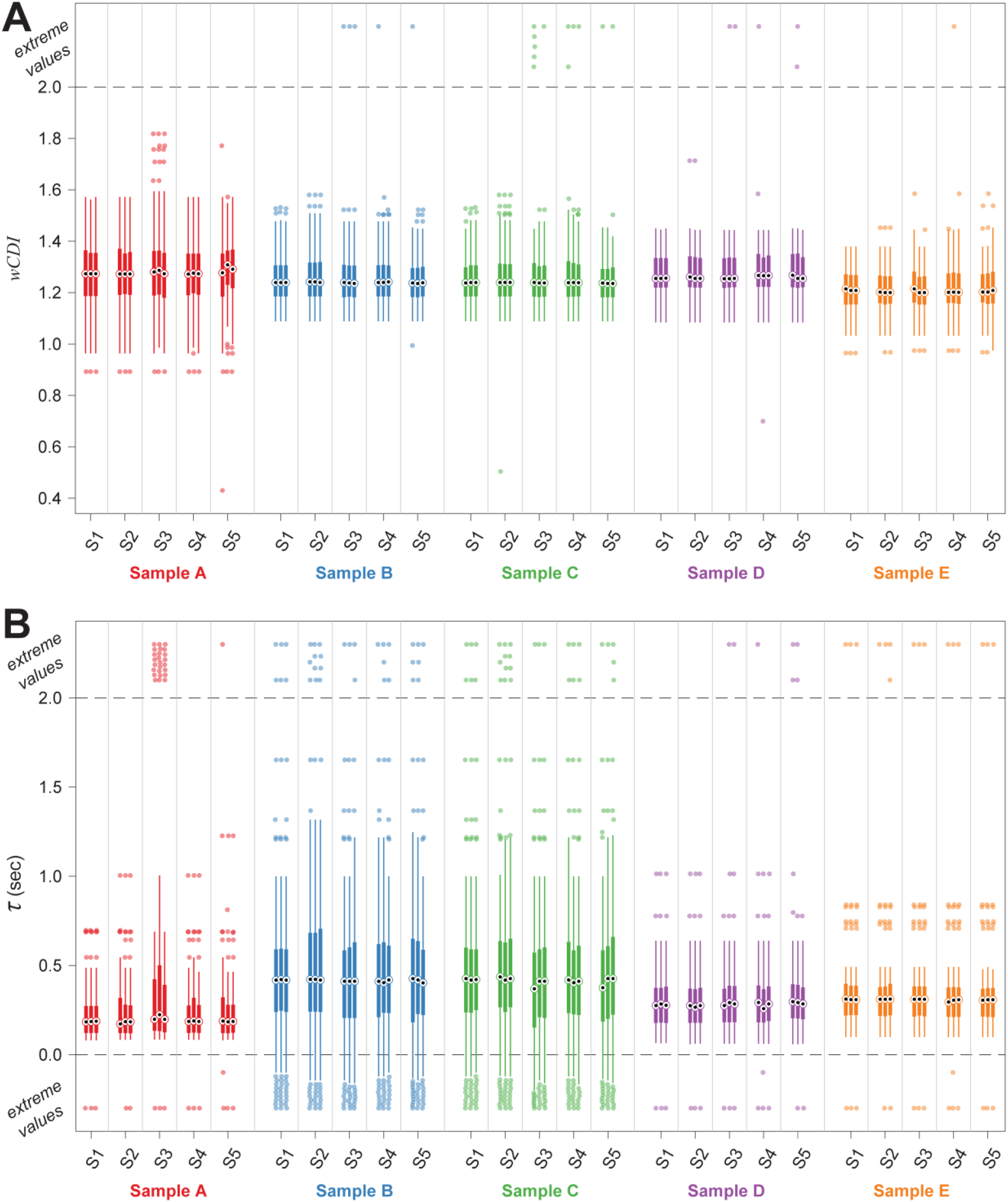
Intra- and inter-user comparison of results obtained using the mechano-NPS data processing pipeline. Processed data obtained from measurements of AP-1060 cells. Data sets A–C were obtained from measuring untreated cells, while data sets D–E were obtained from measuring cells treated with LatrunculinA (LatA). Plots show *wCDI* (A) and recovery time constant *τ* (B) results, respectively. Processed data is shown from each of the five subjects across three replicate data processing tasks. Data is presented as originally returned after the data processing task was completed (i.e., before erroneous measurements were excluded). Sample medians are represented by black dots; sample interquartile ranges are represented by thick lines; outliers are represented by filled circles and are defined as 1.5 times the inter-quartile range. Extreme values are represented as filled circles placed above or below the dashed lines and are defined as >2 or <0 for both *wCDI* and *τ*. Number of cells found in each observation ranged from 49–82.

The data processing software asks for user input at two different stages for each potential cell measurement (Fig 1C): First, the user must decide whether to save the cell measurement or to discard it (in cases where the presented measurement is not a cell, or when cell measurements overlap). If the user decides to save the cell, they are then asked to set two thresholds that affect the cell phenotype measurements that are recorded (see Methods section for additional details on the data processing pipeline). Since these data sets were obtained using the AP-1060 device design, there are two continuously distributed mechanical phenotyping measurements recorded for each cell: *wCDI* and recovery time constant *τ*. The number of cells found by a single subject in a single observation from a single raw data file ranged from 49–82.

We first analyzed the consistency of each user’s decisions to save or discard cell measurements. We calculated percent-agreement and performed Fleiss’s kappa analysis on the decision to save or skip each cell. The intra-user agreement analysis (Fig 4A) shows that all subjects exhibited a high degree of self-consistency in their decisions to save or discard cell measurements. The experienced software users were designated as showing “perfect agreement” according to Landis & Koch’s interpretation of the Fleiss’s kappa values [17], while the novice users demonstrated “substantial” or “moderate” agreement. For all users, the null hypothesis was rejected in Fleiss’s kappa analysis, indicating that the observed agreement was not accidental. The inter-user Fleiss’s kappa analysis (Fig 4B) shows that the overall consistency between all subjects was moderately high (~80% agreement above chance), and “fair agreement” was observed. Pairwise analysis of the agreement between subjects reveals that the experienced users had a high degree of agreement with each other, as well as with one of the novice users (Subject 3). Subjects 1–3 showed “moderate” or “substantial” agreement with each other, while all other subjects showed “fair agreement” with each other. For all comparisons, the null hypothesis was rejected. Importantly, Fleiss may be an overly conservative measure of agreement because 1) it considers the possibility that users may assign labels randomly, which is unlikely to be the case in this data processing task; and 2) this analysis cannot account for agreement on cells that were skipped in all observations, so the probability of encountering a cell that should be saved (and thus, the probability of agreeing to save a cell by chance) is highly overestimated in the Fleiss calculation. For this reason, we have also calculated the raw percent-agreement value, which directly quantifies the percentage of times that the observations agreed on whether to save or skip a cell. For all intra- and inter-user analyses, the raw percent-agreement was ~85% or higher, and the two experienced users plus Subject 3 showed >90% raw agreement among themselves.

**Fig 4.**
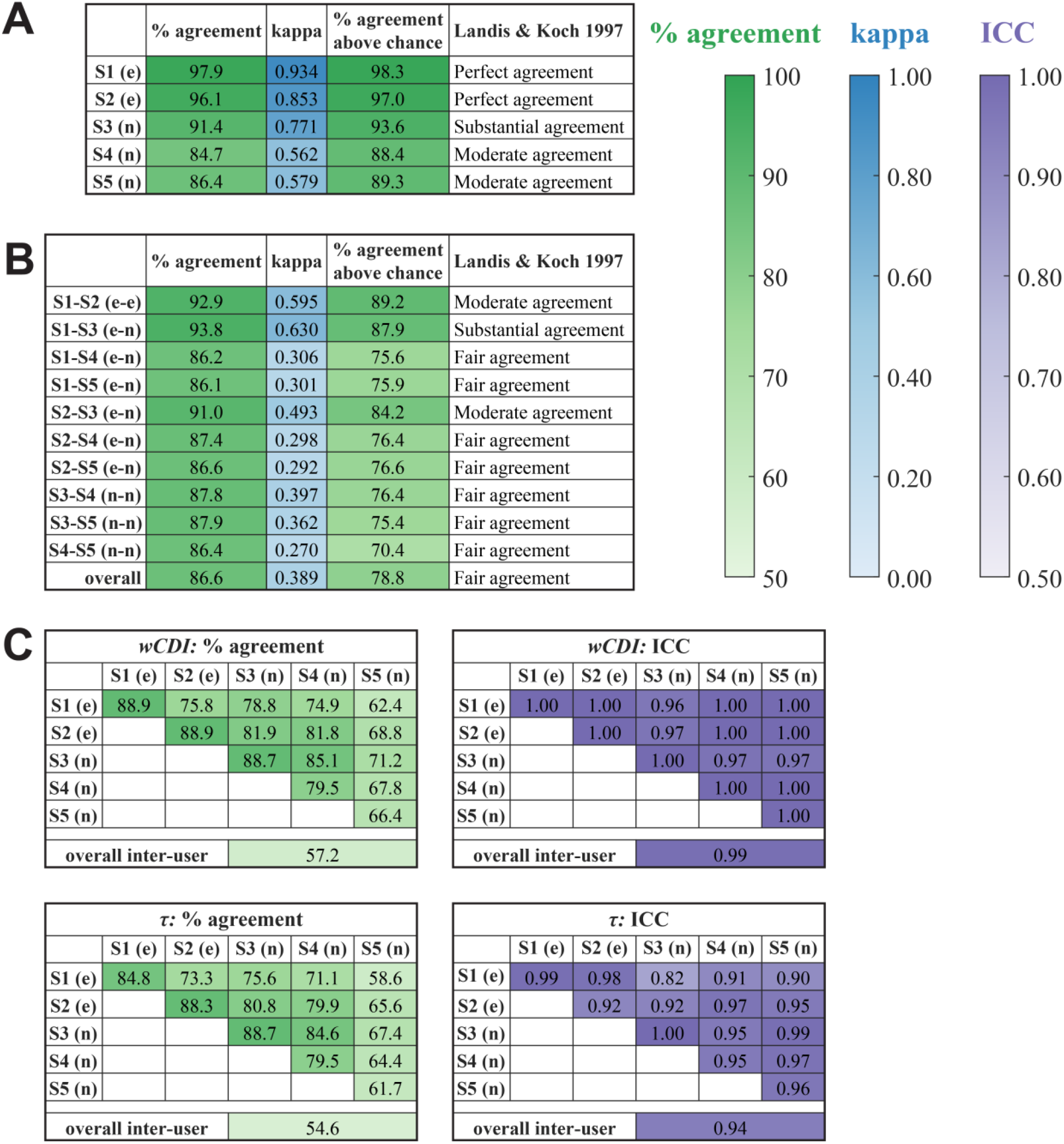
Intra- and inter-user reliability analysis of the mechano-NPS data processing pipeline. Reliability analysis results obtained from the data presented in Fig 3. (e) and (n) designate experienced and novice users, respectively. Number of cells found in each observation ranged from 49–82. Intra- (A) and inter-user (B) agreement analysis examines the consistency of users’ decisions to save or discard cell measurements. “% agreement” quantifies the percentage of potential cell events in which all observations agreed on whether to save or discard the event. Fleiss’s kappa analysis was also performed to determine the “kappa” value as well as the “% agreement above chance.” “Landis & Koch 1997” indicates the interpretation of the kappa value according to [17]. For all comparisons, Fleiss’s kappa analysis rejected the null hypothesis that the observed agreement was accidental (*p <* 10^−10^). (C) Intra- and inter-user concordance analysis examines the consistency of the observed quantitative cell phenotypes for each of the two phenotype variables: *wCDI* and recovery time constant *τ*. “% agreement” quantifies the percentage of cell events in which all observations found the same phenotype value, within tolerance. We also calculated the intra-class correlation (ICC) value, which quantifies the degree of correlation among the observations. For all comparisons, ICC analysis rejected that the null hypothesis that *ICC* = 0 (*p <* 10^−10^).

We then analyzed the consistency of the observed quantitative cell phenotypes, which can be affected by the thresholds set by the user. For each of the two mechanical phenotyping variables, we calculated the percentage of cells that found an equivalent value in each observation, and we also calculated the intra-class correlation (ICC) to quantify the correlation in the measured values between observations. The intra-user analysis (Fig 4C, along the diagonals) shows that all subjects found equivalent values for both variables a majority of the time; most of the users found equivalent values for both variables ~85% of the time or more, although one of the novice users only found equivalent values for the variables ~60–70% of the time. However, the intra-user correlation of values was very high for all subjects (*ICC* > 0.9), demonstrating a very high degree of self-consistency in both variables. The inter-user analysis (Fig 4C) was similar to the intra-user analysis, showing that subjects found equivalent values with each other for both *wCDI* and *τ* a majority of the time. We again find that the two experienced users and one of the novice users (Subject 3) shower a higher degree of consistency among themselves than with the other two novice according to the percent-agreement. However, the ICC analysis showed a very high degree of correlation in both variables among all users (*ICC* > 0.9 in all but one comparison). For all intra- and inter-user analyses, the null hypothesis that *ICC* = 0 was rejected for both *wCDI* and *τ*.

Overall, the agreement and correlation analyses demonstrate that the semi-manual mechano-NPS data processing platform enables reproducible results. Users demonstrate consistent results upon repeated analysis of the same data set, and users find results that are consistent with each other. Although experienced users do show a higher degree of consistency than novice users, novice users of this software are able to achieve reliable results right away. This finding is critical for ensuring that the mechano-NPS platform, and its associated data processing pipeline, is reproducible even when adopted by new and inexperienced users.

### Comparing mechano-NPS results from experiments performed on different instrumentation platforms

Finally, we investigated whether results from a mechano-NPS experiment are reproducible if another research group builds a similar mechano-NPS platform and conducts the same experiment. Two mechano-NPS platforms were used in this comparison: one platform located in Berkeley, California (Site A), and a similar one located in Duarte, California (Site B) (see Methods for a complete list of hardware). Although there were slight differences in the instrumentation used (e.g., digital vs. analog current preamplifier), the platforms are functionally identical. At both sites, an identical mechano-NPS device design was used to measure MCF-10A cells (Fig 1B), and all devices were fabricated at Site A using an identical process (see Methods) [5]. To compare mechanical phenotyping results, replicate vials of cryo-preserved MCF-10A cells were distributed from Site B, thawed, and cultured in identical growth media (see Methods). A researcher at each site measured the cells using that site’s mechano-NPS platform and extracted two mechanical phenotyping parameters—cell deformability and recovery category—as described in previously published work (Fig 1) [5].

Between Site A and Site B results, the MCF-10A mean and median *wCDI* varied by less than 0.4% and 2%, respectively (Fig 5A). Lilliefors tests for *wCDI* distributions determined that both Site A (*p <* 0.001) and Site B (*p <* 0.001) distributions were non-normal, and a Mann-Whitney U-test found that the median *wCDI*s were not statistically significantly different (*p =* 0.055). We then performed a post hoc power analysis of this test to determine the minimum effect size detectable with 80% power and a significance criterion of *α* = 0.05. We computed this effect size to be 0.0275; since the actual effect size for this experiment was only 0.0201, we conclude that the difference in median *wCDI* between Site A and Site B is not a meaningful difference, even though it is statistically significant. We also compared the empirical CDFs for *wCDI* data obtained at Site A and Site B (Fig 5B) and found that they were statistically significantly different (*p =* 0.042), with the maximum absolute deviation occurring between the sample medians.

**Fig 5.**
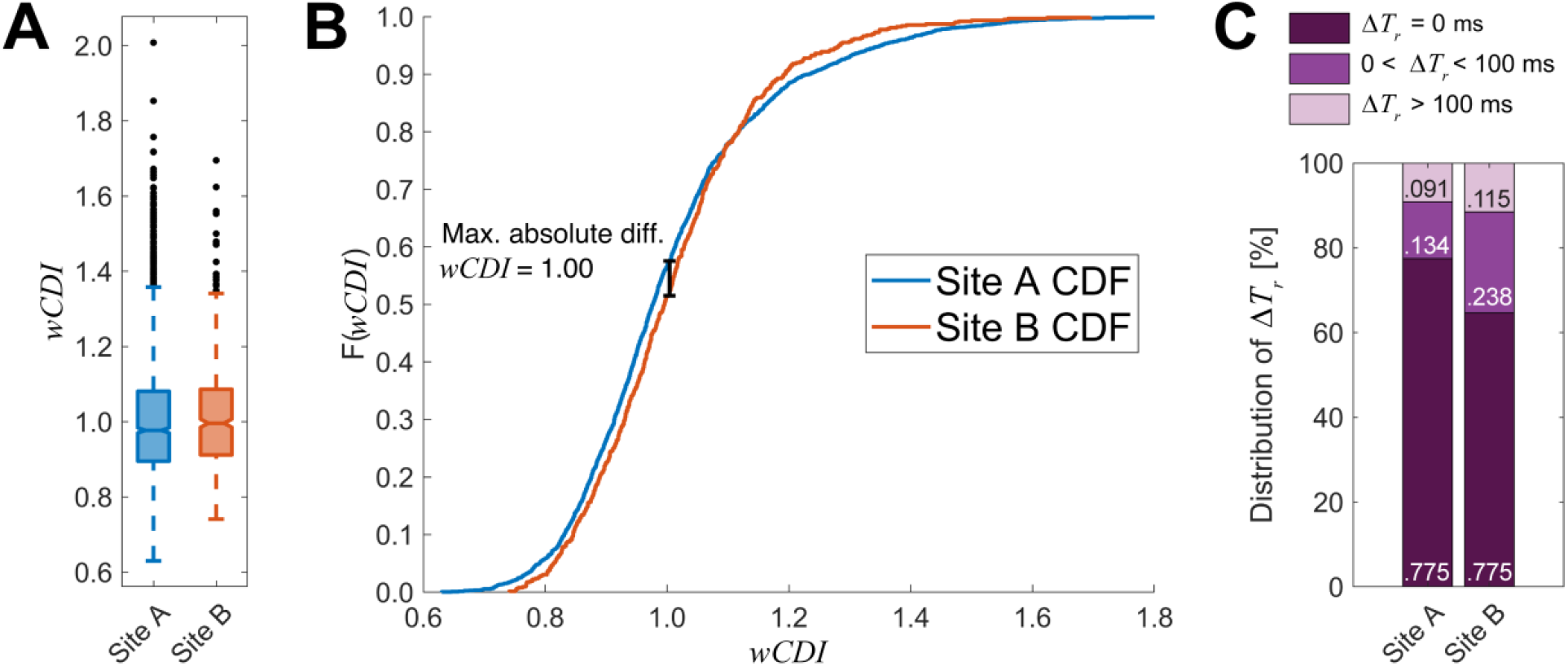
Mechano-NPS results from different instrumentation sets at location Sites A and B are highly reproducible. (A) Box plots of *wCDI* for MCF-10A cells analyzed at Site A and Site B. Black points represent cells with outlier values of *wCDI*, defined as 1.5 times the inter-quartile range. Notches represent 95% confidence intervals for the true sample median for each distribution. (Site A *n =* 1960, Site B *n =* 625). (B) Empirical CDFs for Site A and Site B *wCDI* data. A two-sample Kolmogorov-Smirnov test determined that these data are sampled from different distributions (*p =* 0.042; Site A *n =* 1960, Site B *n =* 625), with the maximum absolute difference occurring at *wCDI =* 1.00. (C) Stacked bar graph representing the relative categorical frequency of MCF-10A cells recovering instantaneously (*ΔT*_*r*_ = 0 ms), within the finite time window (0 < *ΔT*_*r*_ < 100 ms), or failing to recover within the finite time window (*ΔT*_*r*_ > 100 ms). Chi-squared analysis determined that these frequencies were statistically significantly different (*p <* 0.0001; Site A *n* = 1960, Site B *n* = 625; see S5 Table).

We then investigated the reproducibility of MCF-10A recovery categories. Kim et al. classified cell recovery into three categories: immediate, finite, and prolonged. A cell is classified as “immediate” if it is observed to recover its original shape immediately, “finite” if it recovers within a finite time range, or “prolonged” if it does not recover within the finite range (implying slow, prolonged recovery) [5]. We classified the MCF-10A recovery categories measured by Site A and Site B and, using a Pearson’s Chi-squared test, found that the frequencies of recovery categories were significantly different (*p <* 0.0001) with a Cramér’s V of 0.094, which is considered a small-to-medium effect size (Fig 2C) [18–20].

By comparing *wCDI* and recovery category data between the two sites, we observed statistically significant differences according to a standard significance criterion of 0.05. As mechano-NPS is easily capable of measuring hundreds to thousands of cells, it is reasonable that even small differences may be statistically significant. In this case, we would expect differences in cell culturing and handling (e.g., expert vs. novice tissue culture technique, lot-to-lot differences in trypsin-based disassociation solutions, etc.) to account for some differences in measured mechanical properties such as those observed here. Furthermore, based on our characterization of device-to-device variability, we expect that manufacturing variation accounts for the majority of the differences observed between Site A and Site B.

The effect sizes we observed between sites are minuscule in comparison to work by Kim et al. where primary human mammary epithelial cells were immortalized to mimic malignant progression, which we consider biologically relevant and meaningful [5]. Specifically, we measured a between-sites difference in median *wCDI* of 2%, whereas Kim et al. reported differences ranging from 5.8% to 12.8% for their primary strain 122L when treated with shRNA targeting p16 or cyclin D1. For recovery category, we reported a Cramér’s V of 0.094, and while Kim et al. did not report this figure, we computed a value of 0.242 based on their raw data, which is considered medium-to-large [19,20]. Consequently, while our measurements of MCF-10A cells at Site A and Site B produced statistically significantly different results for *wCDI* and recovery category frequencies, the magnitudes of the effect sizes indicate that these differences may not be meaningful.

Based on this experiment, we conclude that the mechano-NPS results from one research group are highly replicable and reproducible when an identical experiment is performed by another group, even when the instrumentation hardware is slightly different.

## Conclusions

Our work characterizing several sources of technical variability and assessing experimental reproducibility represents a novel kind of analysis regarding the use of microfluidic technologies for measuring cell mechanical properties. Increasingly complex and powerful devices can introduce greater opportunity for variability, and the repeatability of the measurements made with these technologies should be addressed accordingly. We investigated three aspects of the reproducibility of an experiment: (1) how technical variation between mechano-NPS devices might affect a measurement, (2) how the results of mechano-NPS measurements might vary due to the researcher processing the data, and (3) how mechano-NPS measurements might vary when performed by different research groups at different location. A study such as this one provides a performance benchmark for other researchers who adopt a technology like mechano-NPS and sets expectations of technical variability to assess the reproducibility of experiments. Especially because single-cell measurements are inherently sensitive to biological heterogeneity, it is important to understand and minimize these sources of technical variability.

## Methods

### Device design

Design parameters for MCF-10A and AP-1060 devices emulated those used as previously published [5,6]. Geometric features were chosen to optimize the signal-to-noise ratio of the microfluidic four-terminal measurement and apply a specific degree of deformation to cells. See Fig 1B for the layout of each device type and Table 1 for each device’s geometric dimensions.

**Table 1.** Geometric dimensions of mechano-NPS devices.

| Microfluidic feature | MCF-10A devices | AP-1060 devices |
| --- | --- | --- |
| Channel height | 22.3 $\mu\text{m}$ | 12.9 $\mu\text{m}$ |
| Inline filter pore width | 22 $\mu\text{m}$ | 20 $\mu\text{m}$ |
| Pore width | 22 $\mu\text{m}$ | 13 $\mu\text{m}$ |
| Pore length | 700 $\mu\text{m}$ | 800 $\mu\text{m}$ |
| Node width | 85 $\mu\text{m}$ | 85 $\mu\text{m}$ |
| Node length | 50 $\mu\text{m}$ | 50 $\mu\text{m}$ |
| Contraction segment width | 10.5 $\mu\text{m}$ | 7.0 $\mu\text{m}$ |
| Contraction segment length | 3000 $\mu\text{m}$ | 2000 $\mu\text{m}$ |
| Recovery segment length | 700 $\mu\text{m}$ | 290 $\mu\text{m}$ |
| Targeted strain in contraction segment | 0.30 | 0.35 |

### Device fabrication

The mechano-NPS channels were fabricated using standard soft lithography. Briefly, a negative-relief master was lithographically fabricated onto a polished silicon substrate using SU-8 epoxy photoresist (MicroChem) (SU-8 3025 for MCF-10A devices, SU-8 3010 for AP-1060 devices.). Polydimethyl siloxane (PDMS) (Sylgard 184, Dow Corning) was mixed at a ratio of 9:1 pre-polymer base to curing agent, degassed with a vacuum desiccator, and then poured onto the negative relief masters. The PDMS was cured by baking at 85°C (358 K) on a hotplate for 2 h, and a PDMS slab containing the embedded microfluidic channel was subsequently excised. The inlet and outlet ports were cored with a biopsy punch (Harris Uni-Core, Fisher Scientific).

Thin-film metal electrodes and contact pads were fabricated on a glass substrate. Briefly, standard photolithography was used to pattern Shipley 1813 photoresist (MicroChem) on the substrate. Electron-gun evaporation was then used to deposit a 75/250/250 Å titanium/platinum/gold thin film onto the patterned substrate, and photoresist liftoff was accomplished with immersion in acetone (JT Baker 9005-05 CMOS grade). For the MCF-10A devices, a gold wet etch solution (GOLD ETCHANT TFA, Transene Company) was drop-cast onto the area of electrodes crossing the microfluidic channel, exposing the Pt electrodes.

Mechano-NPS device fabrication was completed by treating the PDMS slab and glass substrate with pre-fabricated electrodes with oxygen plasma (Harrick Plasma, 450 mTorr (60 Pa), 30 W, 2 min). 20 μL of 2:1 methanol (ACS Grade, VWR BDH1135-4LG) to water (18.2 MΩ) was drop-cast onto the glass substrate to aid in alignment. The slab and substrate were then aligned, mated, and baked on a hotplate to evaporate the methanol-water mixture. The MCF-10A devices were baked at 85°C for 2 h, and the AP-1060 devices were baked at 125°C for 5 min.

### Cell culture

AP-1060 cells (DSMZ ACC 593), a gift from Dr. S. Kogan, University of California, San Francisco, CA, U.S.A., were cultured at 37°C with 5% CO_2_. They were initially seeded at a density of 1 × 10^6^ cells/mL in growth media comprised of 70% Iscove’s modified Dulbecco’s medium (IMDM, Gibco 12440053), 20% fetal bovine serum (FBS, VWR 89510-186), 10% conditioned medium from cell line 5637 (ATCC HTB-9), and 1X Penicillin-Streptomycin (Gibco 15070063). Cells were passaged when suspension cultures reached a density of 2.5 × 10^6^ cells/mL. Conditioned medium from cell line 5637 was prepared by seeding 2.5 × 10^5^ cells in 10 mL of growth media consisting of 90% RPMI-1640 (Corning 10-040-CV), 10% FBS, and 1X Penicillin-Streptomycin. Medium was changed after 24 h and collected after another 24 h. Before adding to AP-1060 growth media, the conditioned medium was filtered using a 0.22 μm polyethylsufone filter (Millipore Sigma SLGPM33RS). For intra- and inter-user repeatability experiments, two samples of AP-1060 cells were treated with LatA (Abcam ab144290). LatA was reconstituted in ACS reagent grade ethyl alcohol (Sigma-Aldrich 459844) to a stock concentration of 2 mM, then aliquoted and kept frozen at –20°C until use. LatA was thawed and added to growth medium at a concentration of 2 μM. Cells were subsequently incubated in the LatA-supplemented growth medium for 30 min at 37°C with 5% CO_2_. After 30 min, LatA-treated cells were collected by centrifuging at 200 RCF for 5 min. Cells were washed once with 1X phosphate buffered saline (PBS) and centrifuged again for 5 min at 200 RCF before immediately being resuspended for mechano-NPS measurements (see below).

MCF-10A cells (ATCC CRL-10317) were cultured at 37°C and 5% CO2 in M87A medium containing cholera toxin and oxytocin at 0.5 ng/mL and 0.1 nM, respectively. Cells were passaged when adherent culture reached 75% confluence. After the fifth passage, cells were frozen at a concentration of 1 × 10^6^ cells/mL for use in reproducibility testing at Sites A and B. Upon thawing at Sites A and B, the cells were seeded at a density of 1 × 10^5^ cells/mL and media was changed every 48h until they were passaged at 75% confluence, with the last media change 24h prior to cell dissociation. Cell dissociation was accomplished by incubating cells in 0.25% trypsin-EDTA (Gibco, 25200056) at 37°C and 5% CO_2_ for 5 min, followed by trypsin neutralization with complete M87A medium (twice the volume of trypsin solution used). MCF-10A cells were collected by centrifuging at 200 RCF for 5 min, washed once with 1X PBS, then centrifuged again for 5 min at 200 RCF before immediately being resuspended for mechano-NPS measurements (see below).

### Mechano-NPS measurements

Suspension-culture AP-1060 cells were prepared for mechano-NPS by first transferring the cell suspension into microcentrifuge tubes and centrifuging for 5 min at 200 RCF. After aspirating the growth-medium supernatant, cells were washed with 1X PBS solution and centrifuged again for 5 min at 200 RCF. The PBS was aspirated, and the AP-1060 pellet was resuspended in 1X PBS supplemented with 2% FBS to reduce adhesion between cells and cell adhesion to PDMS. The cell density was diluted to 3 × 10^5^ cells/mL in 1X PBS supplemented with 2% FBS.

MCF-10A cells were prepared for mechano-NPS in a similar fashion. Cells were dissociated from the culture dish by incubating with 0.05% trypsin-EDTA (Gibco #25300062) for 10 min at 37°C, centrifuged for 5 min at 200 RCF, and resuspended in 1X PBS to a cell density of 1 × 10^5^ cells/mL.

To perform mechano-NPS measurements, diluted cell suspensions were aspirated into polytetrafluoroethylene (PTFE) tubing (1/32” (0.793 mm) ID, 1/16” (1.59 mm) OD, Cole-Parmer EW-06407-41) using a 1 mL slip-tip syringe (BD 309659) fitted with a 20-ga (0.91 mm) blunt-tip needle (Jensen Global JG20-0.5TX). For MCF-10A experiments, the tubing was inserted into a 25.4 mm (length) x 20-ga (0.91 mm, OD) 304 stainless steel connector (New England Small Tube Corp) which was then inserted into the inlet ports. For AP-1060 experiments, the tubing was directly inserted into PDMS inlet ports. The needle was then disconnected from the syringe and connected to an Elveflow OB1 microfluidic pressure controller. A nominal inlet pressure of 200 mbar (20 kPa) for MCF-10A experiments and 80 mbar (8 kPa) for AP-1060 experiments was applied to induce flow through the mechano-NPS device. A four-terminal current measurement was performed as previously described, using a DC potential of 2.5 V for MCF-10A experiments and 3 V for AP-1060 experiments, and measuring the current at a sample rate of 50 kHz [5,7,8,21].

The instrumentation used to acquire the mechano-NPS data consisted of a custom-printed circuit board (PCB) to perform the four-terminal measurement, a benchtop power supply, a current preamplifier, and a PCIe DAQ to interface with a computer. The data was recorded using custom MATLAB software as previously published [5]. The same custom PCB was used at Site A and Site B, and the circuit diagram can be found in Kim et al. [5]. Site A used a Keysight E36311A power supply, a DL Instruments 1211 current preamplifier, and a National Instruments PCIe-6351 DAQ. Site B used a Hewlett-Packard/Agilent E3630A power supply, a Stanford Research Systems SR570 current preamplifier, and a National Instruments PCIe-6351 DAQ. The raw data was processed using custom MATLAB code as previously published to calculate the size and mechanical properties of measured cells [5–8].

### Data processing and analysis for mechano-NPS

Current data sampled at 50 kHz was first low-pass filtered using a 200-sample-wide rectangular moving average filter. The filtered signal was downsampled to 2.5 kHz, then detrended using asymmetric least squares smoothing [22]. The element-wise difference of the entire data vector was computed and thresholded according to user-supplied values; for example, for AP-1060 cell measurements, we used a normalized drop-change in current (relative to baseline) cutoff of 2 × 10^−4^ for pores and 1 × 10^−3^ for contraction segments. As a cell enters or exits a pore, it causes a step change in the current, which manifests as an extreme value in the first-order discrete time-difference in current. Thus, the threshold separates noise-related fluctuations from these step changes in current. As cells of different sizes generate step changes of varying amplitude, the threshold is set to a value specific to the signal-to-noise ratio of the data. By identifying these step changes, pulse and subpulse boundaries are established, allowing for calculation of subpulse amplitude (i.e., cell size) and duration (i.e., cell velocity). Mechanical parameters are computed as previously described [5,6].

The user-dependent data processing pipeline is described in Fig 1C (code available at https://github.com/sohnlab/mechanoNPS_Li-et-al-2020/releases/tag/mNPS_2020). By finding peak locations in the first-order difference in current, a list of potential data “windows” where a pulse that might belong to a cell are generated. Many windows are automatically classified by the software as signal interference and discarded (e.g., if a subpulse is missing from the window). For the remaining windows, manual confirmation regarding whether the pulse belongs to a cell is necessary through a MATLAB CLI. Threshold values are adjusted as needed by the user before confirming to the program to extract information from the pulse in the window. The user can choose to let the program automatically compute threshold values based on the peak heights in the window or to manually adjust these values.

### Statistical methods and analysis

Statistical outliers, defined as more than 3 median absolute deviations from the sample median, were included in all statistical tests. Erroneous measurements were identified based on cell velocity by first excluding statistical outliers and then setting a cutoff of ± 4 median absolute deviations from the sample median velocity in either the reference pore or contraction segment. Erroneous measurements were excluded from all statistical tests and analyses unless otherwise stated.

Data for *wCDI* and recovery time constant were tested for normality using a Lilliefors test implemented in MATLAB R2020a. To evaluate differences in non-normal distributions, a Kruskal-Wallis test for non-parametric analysis of variance (ANOVA) across groups was implemented in MATLAB R2020a. For pairwise comparisons, a Mann-Whitney U-test with a Bonferroni correction (when applicable) was implemented in MATLAB R2020a. To determine if the probability distributions of *wCDI* and recovery time constant were equal, a two-sample Kolmogorov-Smirnov test with a Bonferroni correction (when applicable) was implemented in MATLAB R2020a. For recovery category data, the proportions of cells in each recovery category were analyzed with a Pearson’s Chi-squared test implemented in MATLAB R2020a.

Intra- and inter-user reliability analysis was performed on cell phenotype data resulting from each data processing observation of the same five raw data files. For all analyses of a given comparison, only cells that were saved in any observation within the comparison group were considered. The percent-agreement was calculated according to the decision to save or discard each observed cell, thus quantifying how often the observations agreed on whether to save or discard a cell (erroneous measurements were not excluded from this analysis). Fleiss’s kappa analysis was also performed according to the decision to save or discard each cell (using an implementation by Shah, 2020 in MATLAB R2020a), with the significance criterion of *α =* 0.05 referring to the threshold beyond which the agreement is statistically significantly better than chance (erroneous measurements were not excluded from this analysis) [23, 24]. Confidence intervals and p-values for Fleiss’s kappa analysis are reported in S2 Table. ICC analysis was performed on the data for *wCDI* and recovery time constant using the *irrNA* implementation in RStudio version 1.2 [25,26], using a 2-way mixed-effects model to evaluate single-rater absolute agreement, with the null hypothesis that *ICC =* 0 evaluated at *α =* 0.05. S3 Table reports confidence intervals and p-values for ICC analysis, as well as the effect of excluding erroneous measurements in this analysis. Additionally, percent-agreement was calculated on the data for *wCDI* and recovery time constant, calculating the percent of cell events in which all observations found an equivalent value for the measured phenotype. Measured values were considered equivalent if the difference was within a tolerance of 1 x 10^−10^ multiplied by the minimum absolute value observed for the measured phenotype. S4 Table reports tolerance values used as well as the effect of excluding erroneous measurements on this analysis.

## Acknowledgments

This work was supported by the National Institutes of Health (NIH 1R01EB024989-01 and NIH 1R01CA190843-0). B.L. and K.L.C. were supported by National Science Foundation Graduate Research Fellowships. N.K.L. was supported by the Ralph A. Seban Heat Transfer Fellowship. B.L., K.L.C., N.K.L., and L.L.S. were supported by a generous gift from Michael and Margaret Checca. The authors also thank M. Kozminsky, T.R. Carey, and R. Falcón-Banchs for their insights and discussion of our work.

## Data availability statement

The final data set (comprising four files) is available from the BioStudies database at www.ebi.ac.uk/biostudies/ (accession number S-BSST632).

## Notes

https://www.ebi.ac.uk/biostudies/studies/S-BSST632?query=S-BSST632

https://github.com/sohnlab/mechanoNPS_Li-et-al-2020/releases/tag/mNPS_2020

### Competing Interest Statement

L.L.S. is listed as an inventor on the patent application for "Mechano-node pore sensing" (US201662407425P, WO2018071731A1).

