## Supporting Information for "Evaluating sources of technical variability in the mechano-node-pore sensing pipeline and their effect on the reproducibility of single-cell mechanical phenotyping"

**S1 Table. Lilliefors tests for mechanical phenotyping data to determine distribution normality.**

*wCDI*

| | *p* |
|---|---|
| Device 1 | 0.086 |
| Device 2 | 0.500* |
| Device 3 | 0.016 |

Recovery time constant $\tau$

| | *p* |
|---|---|
| Device 1 | 0.009 |
| Device 2 | 0.126 |
| Device 3 | 0.072 |

The whole-cell deformability index *wCDI* (left) and recovery time constant $\tau$ (right) of AP-1060 cells measured on three different mechano-NPS devices were tested for normality using a Lilliefors test. A p-value less than 0.05 indicates a failure to reject the null hypothesis that the distribution of *wCDI* or recovery time constant for that device came from a normal distribution with an unspecified mean and standard deviation. *The test statistic exceeded the tabulated values in the MATLAB R2020a implementation of the Lilliefors test.

**S2 Table. Fleiss's kappa analysis for the mechano-NPS data processing pipeline.**

| Comparison | Subject(s) | kappa | lower bound | upper bound | *p* |
|---|---|---|---|---|---|
| intra-user | subject1 | 0.934 | 0.919 | 0.949 | 0 |
| | subject2 | 0.853 | 0.837 | 0.868 | 0 |
| | subject3 | 0.771 | 0.756 | 0.786 | 0 |
| | subject4 | 0.562 | 0.547 | 0.578 | 0 |
| | subject5 | 0.579 | 0.564 | 0.594 | 0 |
| inter-user | sub1-sub2 | 0.595 | 0.573 | 0.616 | 0 |
| | sub1-sub3 | 0.630 | 0.610 | 0.651 | 0 |
| | sub1-sub4 | 0.306 | 0.288 | 0.325 | 0 |
| | sub1-sub5 | 0.301 | 0.282 | 0.320 | 0 |
| | sub2-sub3 | 0.493 | 0.474 | 0.513 | 0 |
| | sub2-sub4 | 0.298 | 0.280 | 0.316 | 0 |
| | sub2-sub5 | 0.292 | 0.274 | 0.310 | 0 |
| | sub3-sub4 | 0.397 | 0.380 | 0.415 | 0 |
| | sub3-sub5 | 0.362 | 0.344 | 0.380 | 0 |
| | sub4-sub5 | 0.270 | 0.252 | 0.287 | 1.11e–15 |
| | overall | 0.389 | 0.383 | 0.394 | 0 |

Five subjects analyzed raw mechano-NPS data taken from AP-1060 cells; each subject processed each of the five blinded raw data files three different times using the mechano-NPS data processing software. The resulting list of cell measurements was analyzed using Fleiss's kappa to quantify the inter- and intra-user agreement on whether to save or skip a given cell measurement. The kappa value is reported along with lower and upper bounds for the 95% confidence interval. A p-value less than 0.05 indicates a failure to reject the null hypothesis that the observed agreement is accidental. This analysis was performed on all cell measurements, including those identified as erroneous. Number of cells found in each observation ranged from 49–82.

**S3 Table. Intra-class correlation of cell phenotype values using the mechano-NPS data processing pipeline.**

| | | | ***including all measurements*** | | | | ***excluding erroneous measurements*** | | | |
|---|---|---|---|---|---|---|---|---|---|---|
| | | **Subject(s)** | ***ICC*** | **lower bound** | **upper bound** | ***p*** | ***ICC*** | **lower bound** | **upper bound** | ***p*** |
| *wCDI* | intra-user | subject1 | 1.00 | 1.00 | 1.00 | 0 | 1.00 | 1.00 | 1.00 | 0 |
| | | subject2 | 1.00 | 1.00 | 1.00 | 0 | 1.00 | 1.00 | 1.00 | 0 |
| | | subject3 | 0.46 | 0.40 | 0.52 | 0 | 1.00 | 1.00 | 1.00 | 0 |
| | | subject4 | -0.02 | -0.08 | 0.05 | 0.673 | 1.00 | 1.00 | 1.00 | 0 |
| | | subject5 | 1.00 | 1.00 | 1.00 | 0 | 1.00 | 0.99 | 1.00 | 0 |
| | inter-user | sub1-sub2 | 1.00 | 1.00 | 1.00 | 0 | 1.00 | 1.00 | 1.00 | 0 |
| | | sub1-sub3 | -0.01 | -0.12 | 0.09 | 0.596 | 0.96 | 0.95 | 0.97 | 0 |
| | | sub1-sub4 | 0.16 | 0.05 | 0.26 | 0.002 | 1.00 | 0.97 | 1.00 | 0 |
| | | sub1-sub5 | 0.00 | -0.11 | 0.11 | 0.473 | 1.00 | 1.00 | 1.00 | 0 |
| | | sub2-sub3 | -0.03 | -0.13 | 0.08 | 0.684 | 0.97 | 0.96 | 0.98 | 0 |
| | | sub2-sub4 | 0.18 | 0.08 | 0.28 | 4.20e-4 | 1.00 | 1.00 | 1.00 | 0 |
| | | sub2-sub5 | 0.02 | -0.09 | 0.13 | 0.368 | 1.00 | 1.00 | 1.00 | 0 |
| | | sub3-sub4 | -0.02 | -0.13 | 0.08 | 0.674 | 0.97 | 0.97 | 0.98 | 0 |
| | | sub3-sub5 | 0.70 | 0.64 | 0.75 | 0 | 0.97 | 0.96 | 0.97 | 0 |
| | | sub4-sub5 | 0.01 | -0.10 | 0.11 | 0.459 | 1.00 | 1.00 | 1.00 | 0 |
| | | overall | 0.10 | 0.06 | 0.14 | 1.01e-7 | 0.99 | 0.98 | 0.99 | 0 |
| $\tau$ | intra-user | subject1 | 0.99 | 0.98 | 0.99 | 0 | 0.99 | 0.98 | 0.99 | 0 |
| | | subject2 | 0.92 | 0.90 | 0.93 | 0 | 0.92 | 0.90 | 0.93 | 0 |
| | | subject3 | 1.00 | 1.00 | 1.00 | 0 | 1.00 | 1.00 | 1.00 | 0 |
| | | subject4 | 0.99 | 0.99 | 0.99 | 0 | 0.95 | 0.94 | 0.96 | 0 |
| | | subject5 | 0.96 | 0.95 | 0.97 | 0 | 0.96 | 0.95 | 0.97 | 0 |
| | inter-user | sub1-sub2 | 0.98 | 0.97 | 0.98 | 0 | 0.98 | 0.97 | 0.98 | 0 |
| | | sub1-sub3 | 0.83 | 0.79 | 0.86 | 0 | 0.82 | 0.79 | 0.86 | 0 |
| | | sub1-sub4 | 0.99 | 0.98 | 0.99 | 0 | 0.91 | 0.89 | 0.92 | 0 |
| | | sub1-sub5 | 0.90 | 0.88 | 0.92 | 0 | 0.90 | 0.88 | 0.92 | 0 |
| | | sub2-sub3 | 0.92 | 0.90 | 0.94 | 0 | 0.92 | 0.90 | 0.94 | 0 |
| | | sub2-sub4 | 0.99 | 0.99 | 1.00 | 0 | 0.97 | 0.96 | 0.97 | 0 |
| | | sub2-sub5 | 0.95 | 0.94 | 0.96 | 0 | 0.95 | 0.94 | 0.96 | 0 |
| | | sub3-sub4 | 0.99 | 0.99 | 0.99 | 0 | 0.95 | 0.94 | 0.96 | 0 |
| | | sub3-sub5 | 0.99 | 0.98 | 0.99 | 0 | 0.99 | 0.98 | 0.99 | 0 |
| | | sub4-sub5 | 0.99 | 0.99 | 0.99 | 0 | 0.97 | 0.97 | 0.98 | 0 |
| | | overall | 0.98 | 0.97 | 0.98 | 0 | 0.94 | 0.93 | 0.95 | 0 |

All subjects' resulting measurements of the two cell phenotype values, *wCDI* and recovery time constant $\tau$, were analyzed to quantify the inter- and intra-user consistency of the observed values.

The intra-class correlation value (*ICC*) is reported along with lower and upper bounds for the 95% confidence interval. A p-value less then 0.05 indicates a failure to reject the null hypothesis that $ICC = 0$. This analysis was performed both including and excluding the cell measurements that were identified as erroneous. Number of cells found in each observation ranged from 49–82.

**S4 Table. Percentage of equivalent measured cell phenotype values using the mechano-NPS data processing pipeline.**

| Variable | Tolerance | Comparison | Subject(s) | % agreement, including all measurements | % agreement, excluding erroneous measurements |
|---|---|---|---|---|---|
| *wCDI* | 4.30e–11 | intra-user | subject1 | 88.9 | 88.9 |
| | | | subject2 | 88.6 | 88.9 |
| | | | subject3 | 87.7 | 88.7 |
| | | | subject4 | 77.8 | 79.5 |
| | | | subject5 | 65.1 | 66.4 |
| | | inter-user | sub1-sub2 | 75.5 | 75.8 |
| | | | sub1-sub3 | 78.8 | 78.8 |
| | | | sub1-sub4 | 73.3 | 74.9 |
| | | | sub1-sub5 | 61.1 | 62.4 |
| | | | sub2-sub3 | 81.7 | 81.9 |
| | | | sub2-sub4 | 79.9 | 81.8 |
| | | | sub2-sub5 | 67.3 | 68.8 |
| | | | sub3-sub4 | 82.8 | 85.1 |
| | | | sub3-sub5 | 69.3 | 71.2 |
| | | | sub4-sub5 | 65.1 | 67.8 |
| | | | overall | 54.8 | 57.2 |
| $\tau$ | 5.80e–9 | intra-user | subject1 | 84.8 | 84.8 |
| | | | subject2 | 88.1 | 88.3 |
| | | | subject3 | 87.7 | 88.7 |
| | | | subject4 | 77.8 | 79.5 |
| | | | subject5 | 60.5 | 61.7 |
| | | inter-user | sub1-sub2 | 73.1 | 73.3 |
| | | | sub1-sub3 | 75.6 | 75.6 |
| | | | sub1-sub4 | 69.6 | 71.1 |
| | | | sub1-sub5 | 57.4 | 58.6 |
| | | | sub2-sub3 | 80.6 | 80.8 |
| | | | sub2-sub4 | 78.1 | 79.9 |
| | | | sub2-sub5 | 64.1 | 65.6 |
| | | | sub3-sub4 | 82.2 | 84.6 |
| | | | sub3-sub5 | 65.5 | 67.4 |
| | | | sub4-sub5 | 61.8 | 64.4 |
| | | | overall | 52.3 | 54.6 |

All subjects' resulting measurements of *wCDI* and $\tau$ were analyzed to quantify the percentage of cell events ("% agreement") in which an equivalent phenotype value was found in all observations, within the reported tolerance. This analysis was performed both including and excluding the cell measurements that were identified as erroneous. Number of cells found in each observation ranged from 49–82.

**S5 Table. Frequencies of MCF-10A cell recovery categories measured at Site A and Site B.**

| | | Number of recovered cells (% recovered cells) | |
|---|---|---|---|
| Recovery category | Time range | Site A | Site B |
| Instantaneous | $\Delta T_r = 0$ ms | 1519 (77.5) | 404 (64.7) |
| Finite | $0 < \Delta T_r < 100$ ms | 262 (13.4) | 149 (23.8) |
| Prolonged | $\Delta T_r > 100$ ms | 179 (9.1) | 72 (11.5) |

MCF-10A cells measured with mechano-NPS at Site A and Site B were classified according to whether they recovered from deformation instantaneously ($\Delta T_r = 0$ ms), within a finite time window ($0 < \Delta T_r < 100$ ms), or had prolonged recovery ($\Delta T_r > 100$ ms).
